# Insulin-stimulated adiponectin secretion in pregnancy is mediated by inhibition of adiponectin ubiquitination and degradation and is impaired in obesity

**DOI:** 10.1101/2021.03.10.432857

**Authors:** Irving L. M. H. Aye, Fredrick J. Rosario, Anita Kramer, Oddrun Kristiansen, Trond M. Michelsen, Theresa L. Powell, Thomas Jansson

## Abstract

In pregnancy, adiponectin serves as an endocrine link between maternal adipose tissue, placental function and fetal growth, with low adiponectin promoting placental function and fetal growth. Circulating adiponectin levels are decreased in obese pregnant women and in gestational diabetes, which is believed to contribute to the insulin resistance and increased risk of fetal overgrowth associated with these conditions. However, the molecular mechanisms governing adiponectin secretion from maternal adipose tissues in pregnancy are poorly understood. Using visceral adipose tissue from lean and obese pregnant mice, we show that obesity in pregnancy is associated with adipose tissue inflammation, ER stress, insulin resistance, increased adiponectin ubiquitination and decreased total abundance of adiponectin. Moreover, adiponectin ubiquitination was increased in visceral fat of obese pregnant women as compared to lean pregnant women. We further observed that insulin prevents, whereas ER stress and inflammation promote, adiponectin ubiquitination and degradation in differentiated 3T3-L1 adipocytes. We have identified key molecular pathways regulating adiponectin secretion in pregnancy. This information will help us better understand the mechanisms controlling maternal insulin resistance and fetal growth in pregnancy and may provide a foundation for the development of strategies aimed at improving adiponectin production in pregnant women with obesity or gestational diabetes.

## INTRODUCTION

Almost 2/3 of American women now enter pregnancy overweight (body mass index, BMI=25-29.9 kg/m^2^) or obese (BMI≥ 30) (1,2) and obese women are at high risk to develop a number of pregnancy complications, including gestational diabetes mellitus (GDM) (3). Obese women are also more likely to deliver an infant that is large for gestational age (>90^th^ percentile)(4–6), has increased adiposity (7,8) and is insulin resistant at birth (9). In addition, babies of women with obesity or GDM are at risk for obesity, diabetes and cardiovascular disease later in life (10–12).

Adipose tissue is a metabolically active organ producing adipokines that influence whole-body insulin sensitivity and energy homeostasis. The adipokine adiponectin enhances insulin sensitivity in the adipose tissues, liver and skeletal muscle. Maternal circulating adiponectin levels decrease across gestation, which may contribute to the physiological state of insulin resistance associated with pregnancy (13). Moreover, circulating adiponectin levels are decreased in pregnant women with obesity and GDM (13–15), and low maternal serum adiponectin levels in women are associated with fetal overgrowth (13,14). Studies in pregnant mice with deletion of the adiponectin gene have shown that low maternal adiponectin causes fetal overgrowth (16,17), and normalization of maternal adiponectin levels in obese pregnant mice increases maternal insulin sensitivity and prevents fetal overgrowth by normalizing placental function (18). Moreover, administration of adiponectin to obese pregnant mice to increase maternal circulating adiponectin to levels observed in lean pregnant mice largely prevented the development of cardiac (19) and metabolic disease (20) in adult offspring. This suggests that low maternal adiponectin is mechanistically linked to the adverse placental and fetal outcomes associated with maternal obesity and GDM, and increasing maternal adiponectin levels may serve as an effective intervention strategy to prevent intrauterine transmission of obesity and metabolic disease. However, the molecular mechanisms governing adiponectin secretion from maternal adipose tissues in pregnancy are poorly understood.

Insulin stimulates the secretion of adiponectin in the murine-derived 3T3-L1 adipocytes, in non-pregnant animals and in humans (21–23). Only 50% of the synthesized adiponectin protein is secreted (24,25), suggesting that degradation plays a key role in regulating adiponectin release. Insulin prevents protein degradation in other tissues by inhibiting the ubiquitin-proteasome system (UPS) (26). The UPS is a highly regulated process controlling the degradation of specific proteins following ubiquitination. Previous reports in non-pregnant mice indicate that the UPS regulates adiponectin protein degradation, contributing to the decline in circulating adiponectin levels with obesity (27,28). However, it is unknown if UPS regulates adiponectin degradation in humans. Moreover, the underlying mechanisms regulating adiponectin ubiquitination are unclear. In non-pregnant individuals, obesity causes inflammation and endoplasmic reticulum (ER) stress in the adipose tissue (29), conditions that impair insulin signaling, resulting in the dysregulation of total body glucose and lipid homeostasis (30). However, the impact of obesity on adipose tissue insulin resistance in pregnancy is not known. Herein, we hypothesized that obesity in pregnancy is associated with adipose tissue insulin resistance and increased adiponectin ubiquitination and degradation. Furthermore, we hypothesized that insulin stimulates whereas inflammation/ER stress inhibits adiponectin secretion by regulation of adiponectin ubiquitination and degradation.

## RESEARCH DESIGN and METHODS

### Mouse model of obesity in pregnancy and collection of adipose tissue

All experimental protocols were approved by the Institutional Animal Care and Use Committees of the University of Texas Health Science Center San Antonio and University of Colorado Anschutz Medical Campus. C57BL/6J female mice (proven breeders) were fed a control (C) or a high fat/high sugar (HF/HS) pelleted diet supplemented by ad libitum access to sucrose (20%) solution as previously reported (18,31). Female mice were mated and studied at embryonic day E18. Gonadal (visceral) fat pads were collected; adipose tissue adiponectin levels, and insulin, inflammatory and ER stress signaling were determined in protein extracts using immunoblotting. Adiponectin ubiquitination was assessed by immunoblotting following immunoprecipitation, as described below.

### Serum and adipose tissue samples from pregnant women

Healthy term pregnant women undergoing planned cesarean section at Rikshospitalet, Oslo, Norway, were enrolled after written informed consent. The study was approved by the Institutional Review Board and the Regional Committee for Medical and Health Research Ethics, South East Norway (reference numbers 2419/2011 and 13885). Maternal blood samples were collected during Cesarean section for serum analysis as previously described (32). Clinical characteristics and serum results are provided in Table 1. Pre-labor cesarean delivery was performed under spinal anesthesia, and omental (visceral) adipose tissue was sampled from a subset of this cohort, immediately after delivery in women with normal pre-pregnancy BMI (N=4, mean 22.35, SD 1.15) and obese women (mean BMI 32.78, SD 2.21). Adipose tissue samples were snap frozen and stored at -80**°**C prior to cryosectioning for proximity ligation assays and protein extraction for immunoblotting.

**Table 1.** Clinical characteristics of the study subjects.

| | Normal BMI<br>Mean ± SD, (%) or<br>Median (Q<sub>1</sub> - Q<sub>3</sub>) | Obese BMI<br>Mean ± SD, (%)<br>Median (Q<sub>1</sub> - Q<sub>3</sub>) | P-value |
| --- | --- | --- | --- |
| N | 14 | 12 |  |
| Age, yrs | 34.6 ± 4.0 | 34.3 ± 4.9 | 0.855<sup>1</sup> |
| Pre pregnancy BMI, kg/m<sup>2</sup> | 22.5 (21.9 – 23.3) | 32.3 (31.6 – 35.8) |  |
| Systolic blood pressure first trimester, mmHg | 111.9 ± 13.4 | 112.0 ± 9.6 | 0.988<sup>1</sup> |
| Diastolic blood pressure first trimester, mmHg | 65.5 ± 9.1 | 71.6 ± 7.3 | 0.074<sup>1</sup> |
| Birthweight, grams | 3780 ± 417 | 3866 ± 521 | 0.656<sup>1</sup> |
| Placenta weight, grams | 613 ± 100 | 659 ± 110 | 0.269<sup>1</sup> |
| Higher education (>12 yrs) | 11 (78.6%) | 8 (66.7%) |  |
| Infant sex male | 9 (64.3%) | 6 (50%) |  |
| Married or partner | 14 (100%) | 12 (100%) |  |
| Pre-gestational diabetes | 0 (0%) | 0 (0%) |  |
| Gestational diabetes | 0 (0%) | 1 (8.3%) |  |
| Gestational age, wks | 39.6 ± 1.0 | 39.0 ± 1.1 | |
| HMW Adiponectin | 3.65 (2.44 – 5.42) | 4.83 ± (3.18 – 5.37) | 0.274<sup>2</sup> |
| Total Adiponectin | 6.82 (4.94 – 10.02) | 9.74 (7.30 – 11.36) | 0.118<sup>2</sup> |
| HMW/total adiponectin ratio | 0.53 ± 0.07 | 0.49 ± 0.05 | 0.0992<sup>1</sup> |
| Glucose, mmol/L | 4.29 ± 0.11 | 4.82 ± 0.19 | 0.473<sup>1</sup> |
| Cholesterol, mmol/L | 5.96 ± 0.21 | 5.98 ± 0.28 | 0.982<sup>1</sup> |
| Insulin, pmol/L | 39.2 (31.7 – 67.1) | 95.9 (60.7 – 116.6) | 0.002<sup>2</sup> |

Statistical significance determined by <sup>1</sup>T-test (2-tailed) or <sup>2</sup>Mann-Whitney U-test (Exact Sig. 2-tailed)

### Proximity ligation assay (PLA) and confocal microscopy

Adipose tissue collected from pregnant women was serially cryosectioned at 4 µm and fixed in 4% paraformaldehyde. Subsequently, sections were blocked using 5% newborn calf serum (NCS) in PBS for 1 hour followed by incubation in anti-adiponectin (anti-rabbit) and anti-ubiquitin (anti-mouse) antibodies for 2 hours at 37° C. PLA probes anti-rabbit PLUS and anti-mouse MINUS were diluted in Duolink dilution buffer and incubated in a pre-heated humidity chamber for 30 min. This was followed by ligation, amplification, and detection according to the Duolink *In Situ* Orange kit (Sigma-Aldrich) manufacturer’s protocol. Confocal microscopy was performed using a Zeiss LSM 780 microscope at 63x magnification using oil immersion. Images were captured in the same laser settings with four Z-step of 0.4 um. The Image J (NIH, USA) was used for quantification of the detected PLA signals.

### 3T3-L1 cell culture and differentiation into adipocytes and treatments

3T3-L1 pre-adipocytes (American Tissue Culture Collection, Rockville, MD) were maintained in Pre-adipocyte Expansion Medium (PEM, 90% Dulbecco’s Modified Eagle Medium [DMEM], 10% newborn calf serum) and split at 70% cell confluency. Differentiation into adipocytes was performed according to standard protocols (33). In brief, 3T3-L1 cells were grown to confluence and maintained for an additional 48hr in PEM (differentiation day 0). Cells were then grown in differentiation medium (90% DMEM, 10% fetal bovine serum [FBS], 1µM dexamethasone, 0.5mM methylisobutylxanthine and 1µg/ml insulin) for 3 days. Subsequently, culture media was replaced with adipocyte maintenance medium (90% DMEM, 10% FBS, 1µg/ml insulin), which was replaced every 2 days until adipocytes had formed (differentiation day 7). Mature adipocytes were identified by lipid droplet formation and the accumulation of neutral lipid dye Oil Red O.

Following 3T3-L1 adipocyte differentiation, culture media was replaced with DMEM, 10% FBS without insulin. 3T3-L1 adipocytes were then incubated with TNF-α or thapsigargin (Tg) for 24h with or without insulin for 3h. Protein lysates were then collected for adiponectin immunoprecipitation and/or immunoblotting analyses as described below.

### Cycloheximide-chase assay of adiponectin degradation

Adiponectin protein half-life was assessed in differentiated 3T3-L1 adipocytes by cycloheximide (CHX)-chase assay as described previously (34). The CHX-chase assay allows for the analysis of protein degradation over time. In brief, 3T3-L1 adipocytes were treated with cycloheximide (an inhibitor of protein synthesis). Protein lysates were collected immediately and subsequently at specific time points indicated in the figures following the addition of the compound. Protein degradation was assessed by immunoblotting analyses as described below.

### Immunoblotting analyses

3T3-L1 adipocytes were harvested, and mouse and human adipose tissue were homogenized in radioimmunoprecipitation (RIPA) buffer (50mM Tris HCl, pH=7.4; 150mM NaCl; 0.1% SDS; 0.5% Na-deoxycholate and 1% Triton X100) containing protease inhibitors and phosphatase inhibitor cocktail 1 and 2 (1:100, Sigma). Immunoblotting was then carried out as described (35). The expression of β-actin was used to control for any differences in loading and transfer.

### Adiponectin and ubiquitin reciprocal immunoprecipitation

Reciprocal immunoprecipitation of adiponectin and ubiquitin was performed using the Pierce™ Co-Immunoprecipitation Kit (ThermoFisher Scientific) with the following modifications. Twenty-five μg of anti-monoclonal adiponectin antibody (Merck Millipore) was immobilized onto coupling resin by incubation overnight at 4^0^C with end-over-end rotation. Immobilized antibodies were then incubated with 1mg of pre-cleared total protein lysate overnight at 4^0^C on a rotator. Subsequently, immunoprecipitated products were eluted and denatured prior to immunoblotting analyses of ubiquitin as described above. To further confirm adiponectin-ubiquitin protein binding, reciprocal immunoprecipitations were performed using ubiquitin antibody followed by immunoblotting of adiponectin.

### Statistics

Unless otherwise stated, data are presented as means ± SEM. Data were analyzed with GraphPad Prism software (GraphPad Software, San Diego, CA). Statistical differences between groups were assessed using students t-test or ANOVA with Dunnett’s test, as appropriate. A P value <0.05 was considered significant.

## RESULTS

### Decreased total adiponectin and increased adiponectin ubiquitination, insulin resistance, inflammation and ER stress in adipose tissue of obese pregnant mice

We previously demonstrated that the obesogenic diet used in this study increases maternal fat mass by 2.2-fold and results in glucose intolerance with normal fasting glucose in pregnancy (31). Maternal circulating insulin, leptin, and cholesterol are increased in obese dams, whereas total and high molecular weight (HMW) adiponectin are decreased (31). Thus, this animal model is associated with maternal metabolic alterations similar to that observed in pregnant women with increased BMI and without gestational diabetes. Importantly, this model of obesity in pregnancy results in increased fetal weight (+18%), replicating fetal overgrowth, which is common in obese pregnant women (4–6,8). Consistent with our previous reports, total adiponectin protein was decreased (**Fig 1A**), and adiponectin ubiquitination was increased (**Fig 1B**) in gonadal adipose tissue of obese (OB) pregnant (E18) mice compared to normal weight pregnant mice (C). These findings show that the previously reported decrease in circulating adiponectin in obese pregnant mice (18,31) is associated with increased adiponectin ubiquitination in adipose tissue. Moreover, we explored whether maternal obesity is associated with adipose inflammation and ER stress, which are known mediators of adipose insulin resistance. OB dams exhibited adipose tissue insulin resistance, as indicated by increased inhibitory serine phosphorylation of IRS-1 (S307) and decreased Akt (T308) phosphorylation (**Fig 1C)**. Obesity was also associated with increased adipose tissue inflammation, as evidenced by decreased IκBα (inhibitor of NF-κB) expression and increased phosphorylation of JNK (**Fig 1C**). Moreover, the adipose tissue ER stress pathways IRE1α and PERK were activated in obese pregnant mice, as demonstrated by their phosphorylation. Activation of ER stress was further supported by increased phosphorylation of eIF2α, a key downstream target of IRE1α and PERK, and increased expression of spliced XBP1 (XBP1s) and BiP (**Fig 1C**).

**Figure 1.**
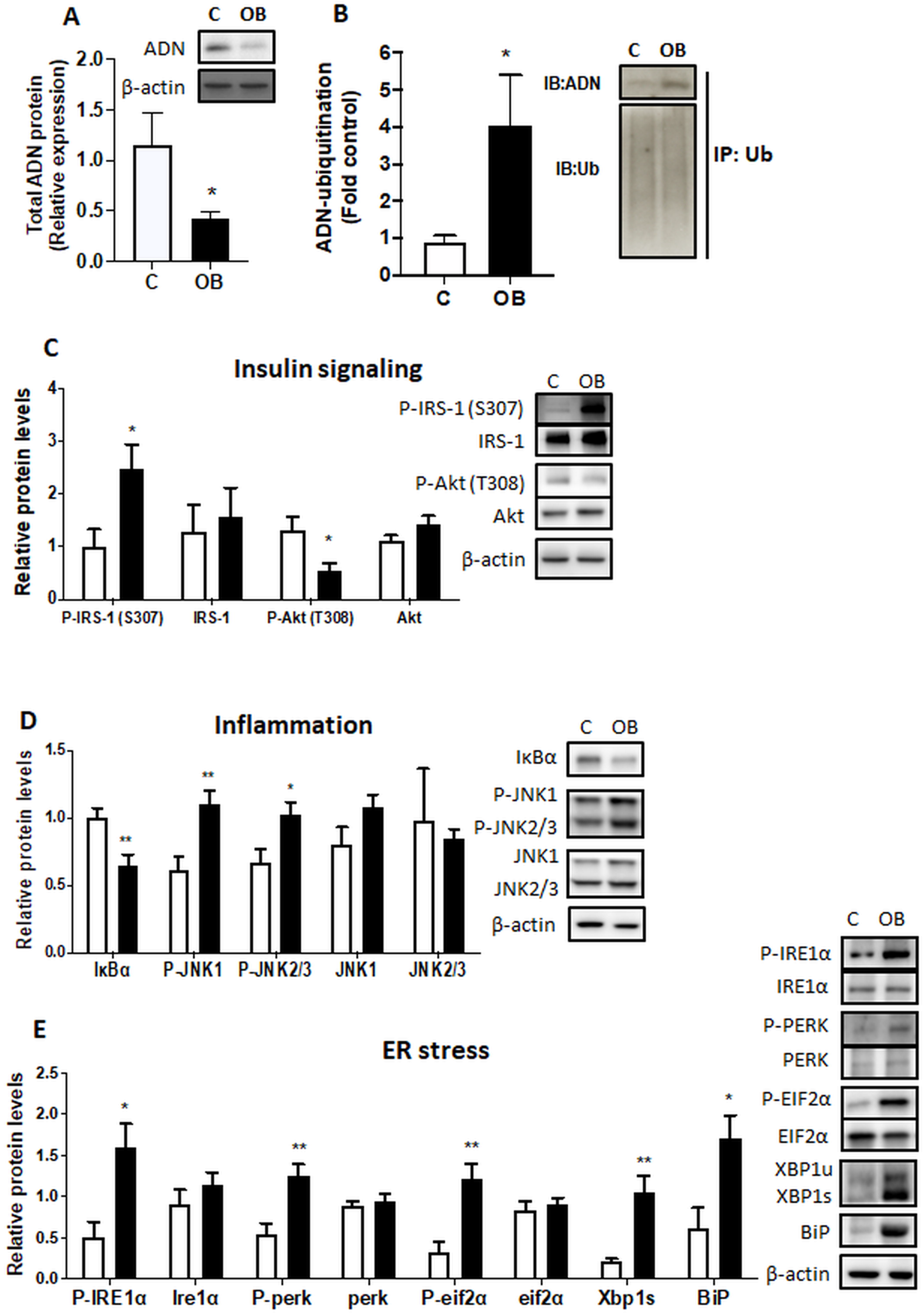
Impact of maternal obesity in mice on adipose tissue adiponectin expression and ubiquitination, insulin signaling, inflammation and ER stress. Gonadal adipose tissue was collected at E18 in pregnant obese [OB] and control [C] mice, and total adiponectin protein expression measured (A). (B) Adiponectin ubiquitination was determined in gonadal adipose tissue by immunoprecipitation (IP) of ubiquitin followed by immunoblotting of adiponectin and ubiquitin in obese pregnant mice. OB dams exhibited adipose tissue insulin resistance (C) as indicated by increased inhibitory serine phosphorylation of IRS-1 (S307) and decreased Akt (T308) phosphorylation. Obesity was associated with increased AT inflammation and ER stress. Adipose tissue inflammation was evidenced by decreased IκBα (inhibitor of NF-κB) expression and increased phosphorylation of JNK. Activation of ER stress pathways was indicated by phosphorylation of IRE1α and PERK, and their downstream target EIF2α, and increased expression of spliced XBP1 (XBP1s) and BiP. Asterisks adjacent to blots indicate significant differences between C and OB. Mean+SEM, unpaired Student’s t test. Figures are representative of n=7 C, n=10 OB. *P<0.05, **P<0.01. C=control, OB=Obese.

### Increased adipose adiponectin ubiquitination in obese pregnant women is correlated with reduced circulating total adiponectin

Using Proximity Ligation Assay in omental tissue samples from pregnant women with normal BMI (range: 21.0–23.4) or obese BMI (range: 30.4–35.2), we observed an increase in adiponectin ubiquitination in obese pregnant women (**Fig 2A, B**), but no changes in total adiponectin in maternal adipose tissues (**Fig 2C**). Similarly, total serum adiponectin levels were not significantly altered by maternal obesity (**Table 1**). However, adiponectin ubiquitination in maternal adipose tissues was inversely associated with maternal serum levels of total adiponectin **(Fig 2D)**.

**Figure 2.**
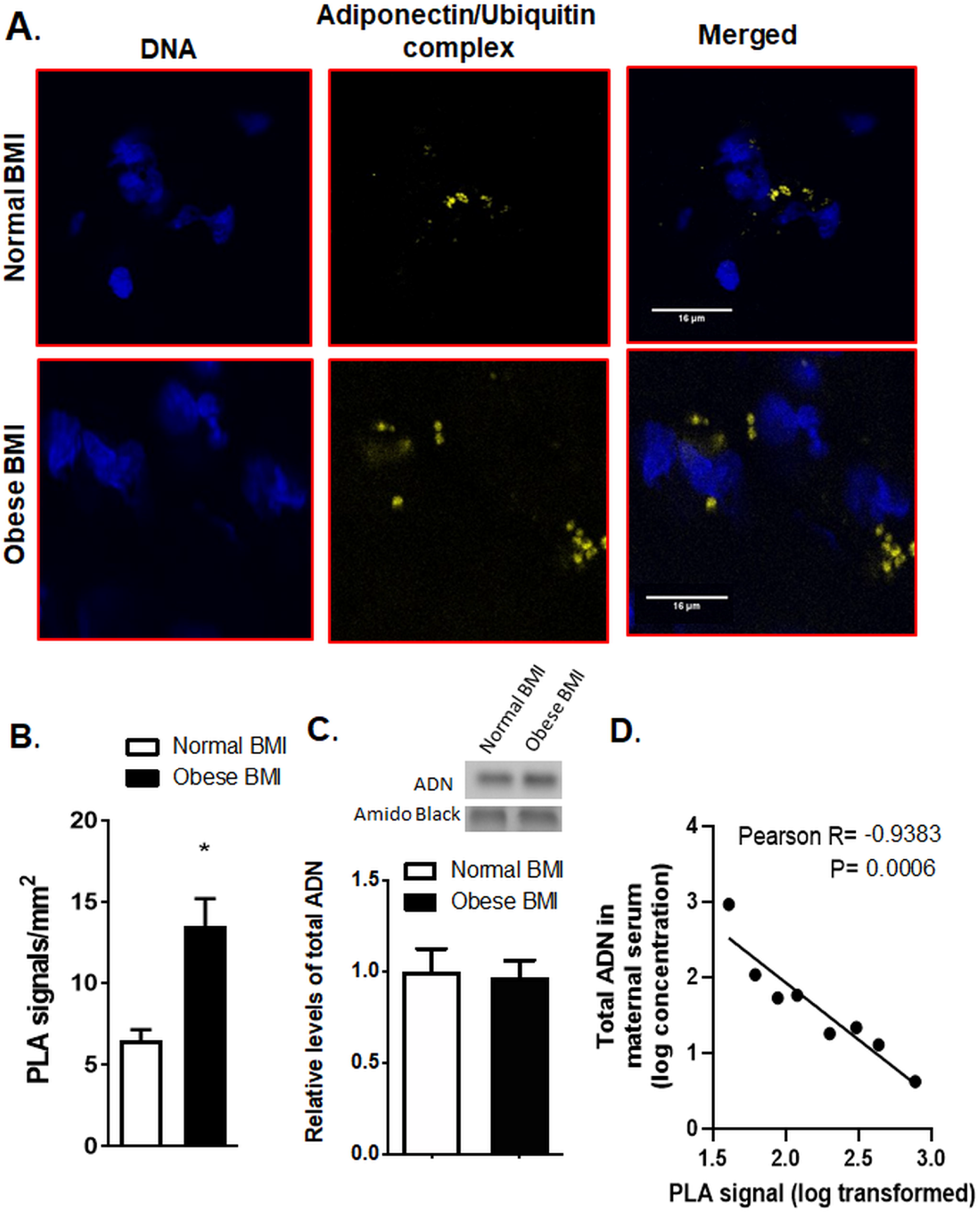
Obesity is associated with increased adipose tissue adiponectin ubiquitination in women. Proximity ligation assay and confocal microscopy were used to determine the association between adiponectin and ubiquitin in the adipose tissue of obese and normal BMI pregnant women (A). In normal BMI, adipose tissue (A, top) adiponectin and ubiquitin associations were limited as evidenced by a low number of PLA signals (yellow dots), suggesting low adiponectin ubiquitination. In contrast, in obese pregnant women (A, bottom), the number of PLA signals was greater, suggesting increased adiponectin ubiquitination as compared to adipose tissue in normal BMI women. Blue = nuclei; Scale bar-16 µm. Bar graph (B) summarizes the proximity ligation assay data of adipose tissue adiponectin-ubiquitin interactions. In each section, at least ten randomly selected microscopic fields were used to calculate the number of adiponectin-ubiquitin association positive sites (yellow dots) per mm^2^. Data were averaged to represent a single adipose tissue. Values are given as mean+SEM; *P<0.05 vs. control; unpaired Student’s t-test; n = 4 adipose tissue/each group. (C) Total adiponectin abundance in the adipose tissue of obese and normal BMI pregnant women as determined by immunoblotting. The bar graph summarizes the Western blotting data. Equal protein loading was assessed by amido black staining. After normalization to amido black, the mean density of control samples was assigned an arbitrary value of 1. Means+SEM, unpaired Student’s t test. (D) Inverse correlation between maternal serum total adiponectin and adipose tissue adiponectin ubiquitination. Linear regression analyses were performed on log-transformed maternal serum concentrations of total adiponectin and adipose tissue PLA signals.

### Insulin increases cellular adiponectin protein abundance by decreasing adiponectin ubiquitination

Using differentiated 3T3-L1 adipocytes as our adipocyte model, we demonstrate that insulin increased cellular adiponectin protein abundance (**Fig 3A)** and secretion to the media (**Fig 3B**). 3T3-L1 adipocytes were treated with insulin followed by CHX-chase assay. Because treatment with CHX markedly reduced cellular adiponectin protein after 3h (**Fig 3C**), demonstrating effective inhibition of protein synthesis, the 3-hour time point was used in subsequent studies. CHX treatment decreased cellular adiponectin protein by >50% after 3h (**Fig 3D**), consistent with previous reports estimating adiponectin protein half-life (24). The degradation of adiponectin was prevented, in part, by stimulation of cells with insulin prior to CHX treatment (**Fig 3D**). Similarly, treatment with a proteasome inhibitor (MG132) counteracted the decline in adiponectin levels, suggesting that proteasome degradation accounts for much of the loss of cellular adiponectin in normally functioning adipocytes. Because UPS is a major mechanism regulating protein degradation, we determined whether adiponectin protein was ubiquitinated and the role of insulin in this process. As shown in **Fig 3E**, ubiquitination of adiponectin was significantly reduced following insulin treatment.

**Figure 3.**
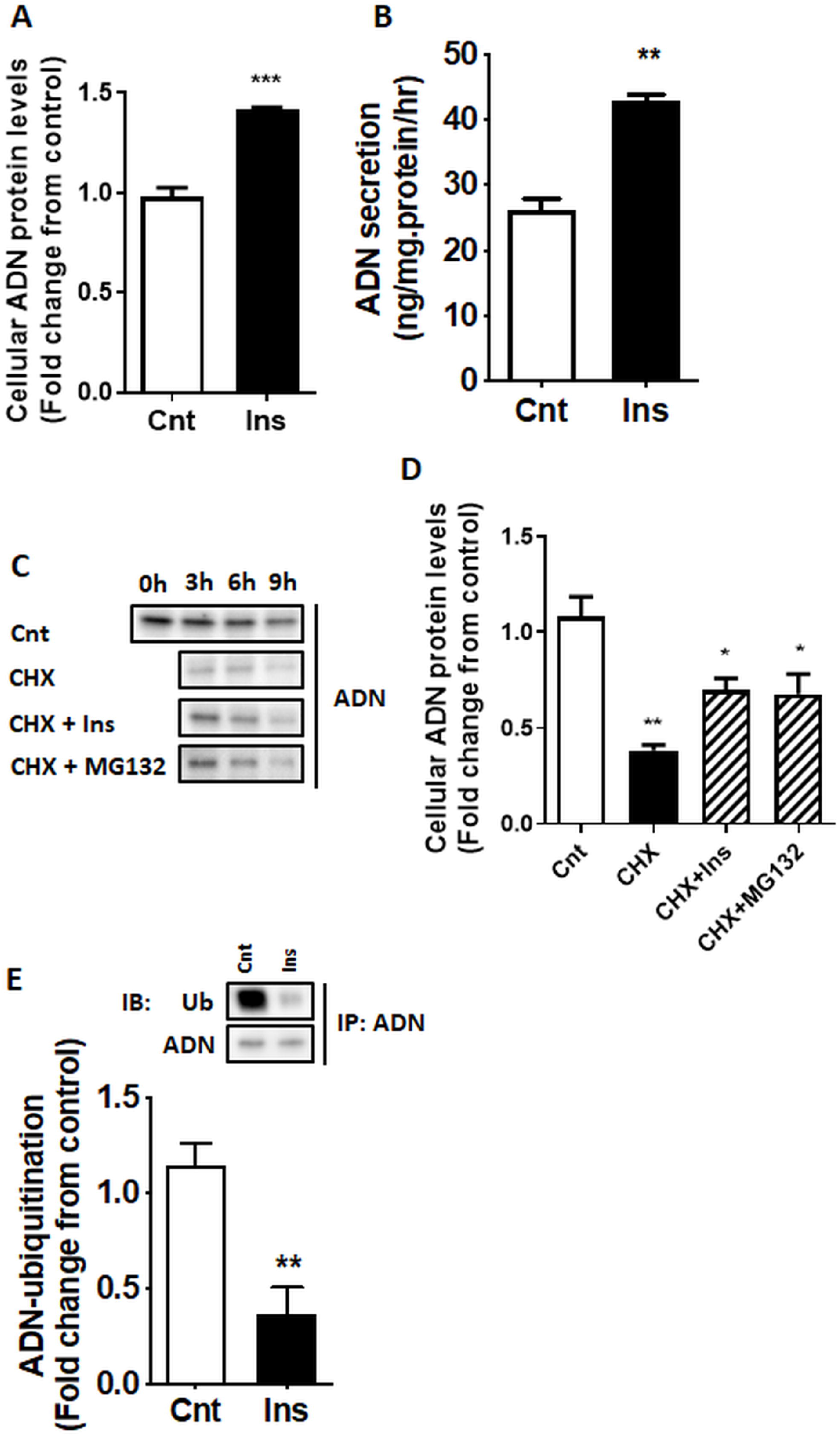
Insulin increases adiponectin secretion and inhibits UPS-mediated degradation of adiponectin. 3T3-L1 cells were differentiated into mature adipocytes using standard protocols. 3T3-L1 adipocytes were cultured for 3h with/without CHX (10μg/ml) or MG132 (10μM), prior to insulin stimulation (10nM). Cellular adiponectin protein expression (**A**) and secretion into culture media (**B**) were determined by immunoblotting analysis and ELISA, respectively. (**C**) Immunoblots showing the time course of adiponectin protein degradation following CHX treatment with/without insulin or the proteasome inhibitor (MG132), and (**D**) the quantification of the results from the 3h time point. (**E**) Ubiquitination of adiponectin after 3h insulin stimulation was determined by immunoprecipitation (IP) of adiponectin followed by immunoblotting (IB) analysis of adiponectin and ubiquitin in the immunoprecipitated protein lysates. Values are Mean+SEM (n=6). Unpaired Student’s t test or One-Way ANOVA *P<0.05, **P<0.01, ***P<0.001. Cnt, Control; Ins, Insulin; CHX, Cyclohexamide.

### Inflammation and ER stress inhibits the effects of insulin on adiponectin abundance, degradation and ubiquitination

Both TNFα (pro-inflammatory cytokine) and Thapsigargin (Tg, ER stress inducer) decreased cellular adiponectin protein levels in 3T3-L1 adipocytes and prevented insulin-stimulated adiponectin production (**Fig 4A**). Furthermore, Tg increased the ubiquitination of adiponectin, whereas both TNFα and Tg prevented the insulin-mediated decrease in adiponectin ubiquitination (**Fig 4B**). CHX-chase assays indicated that TNFα and Tg increased adiponectin protein degradation and prevented the effect of insulin on adiponectin degradation (**Fig 4C**).

**Figure 4.**
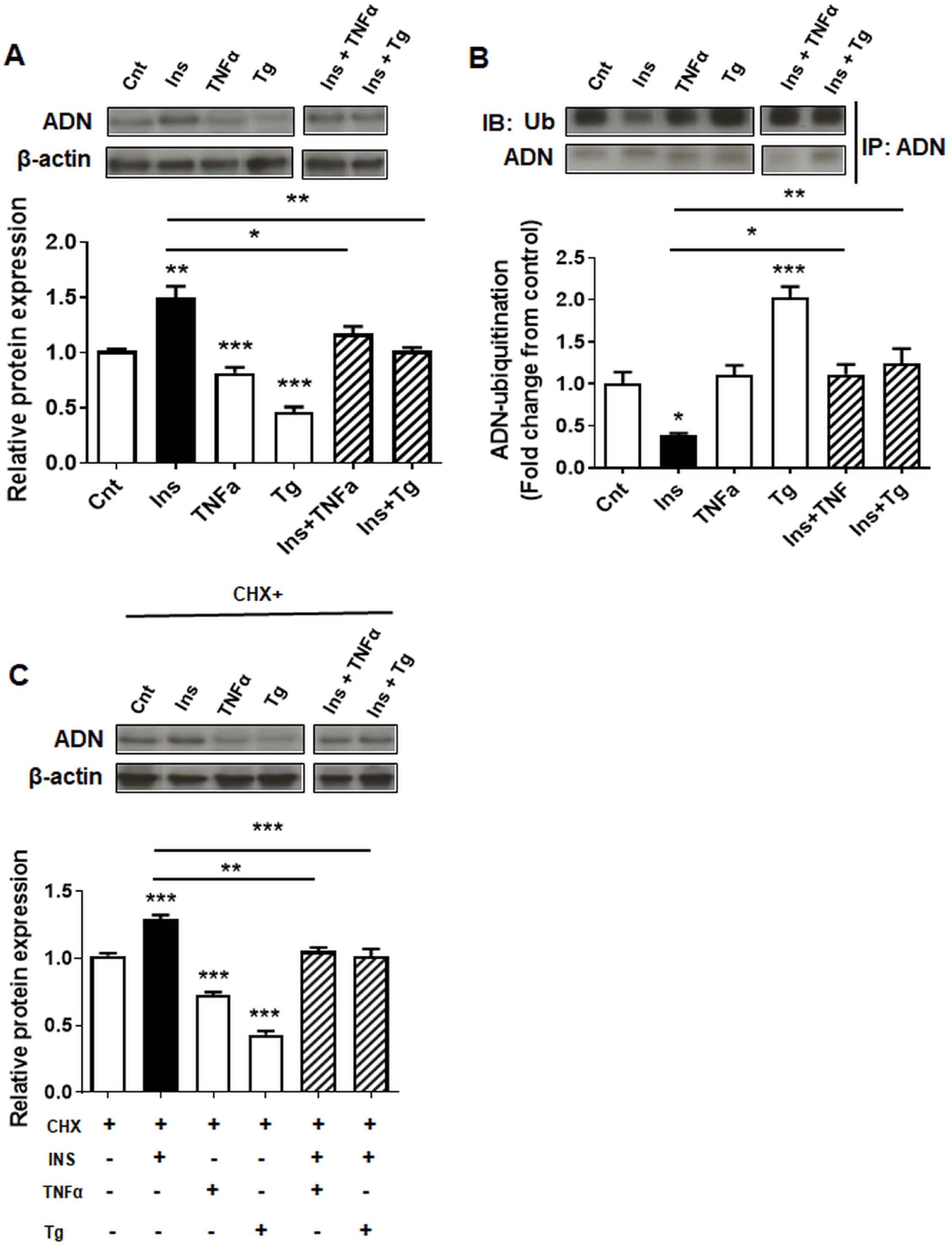
The effects of inflammation and ER stress on insulin-dependent adiponectin protein expression, ubiquitination and degradation. 3T3-L1 adipocytes were treated with the pro-inflammatory cytokine TNFα (20ng/ml) or ER stress-inducing agent Thapsigargin (Tg, 1μM) for 24h prior to stimulation with insulin and/or CHX for 3h. (**A**) Adiponectin protein expression, (**B**) CHX-mediated adiponectin degradation, and (**C**) Ubiquitination of adiponectin. Values are Mean+SEM, n=6. One-Way ANOVA *P<0.05, **P<0.01.

## DISCUSSION

We have identified a molecular mechanism regulating adiponectin secretion, which involves insulin signaling and ubiquitination. This new information will help us better understand the mechanisms underlying the decrease in circulating maternal adiponectin in obesity in pregnancy. Specifically, using visceral adipose tissue from lean and obese pregnant mice, we show that obesity in pregnancy is associated with adipose tissue inflammation, ER stress, insulin resistance, increased adiponectin ubiquitination and decreased total abundance of adiponectin. In addition, as compared to lean pregnant women, adiponectin ubiquitination was increased in visceral fat of obese pregnant women. Moreover, we report that insulin prevents, whereas ER stress and inflammation promote, adiponectin ubiquitination and degradation in cultured adipocytes, which may explain the underlying mechanisms by which obesity mediated adiponectin ubiquitination and degradation contributes to the decline in circulating adiponectin levels in pregnant women (**Fig 5**).

**Figure 5.**
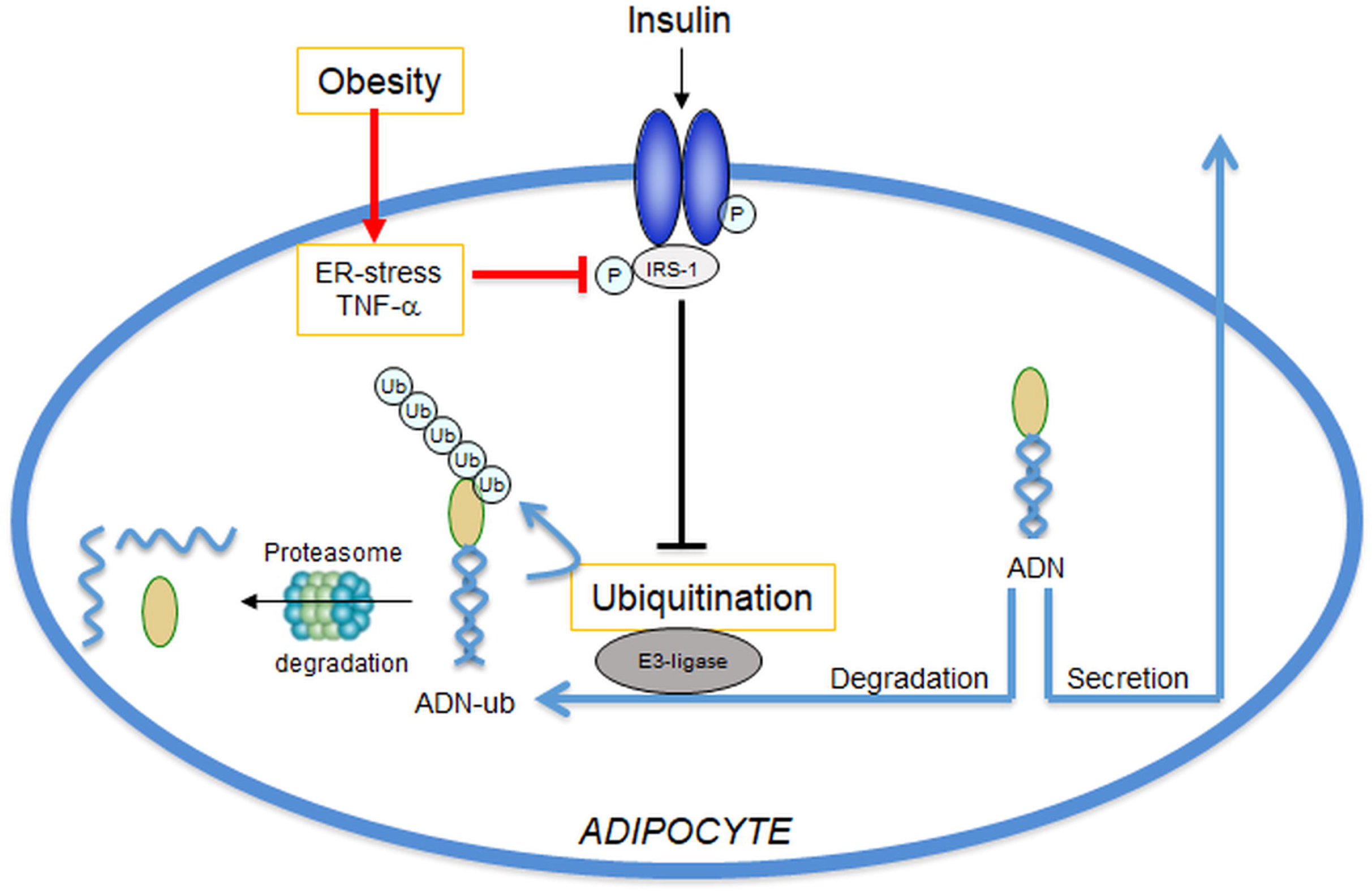
Model for regulation of adiponectin secretion in pregnancy. We propose that insulin stimulates adiponectin secretion in pregnancy mediated by inhibition of adiponectin ubiquitination and degradation, and that obesity is associated with adipose Inflammation, ER stress and insulin resistance resulting in increased adiponectin ubiquitination and degradation.

Circulating adiponectin levels are decreased in pregnant women with obesity (15,37–39) and GDM (16,39), i.e., pregnancy complications associated with glucose intolerance. Adiponectin is well-established to promote glucose tolerance. Therefore, these observations are consistent with the possibility that low adiponectin levels directly contribute to glucose intolerance (40,41). Consistent with this hypothesis, elegant studies in genetically modified mice demonstrate that maternal adiponectin is required to expand beta-cell mass in pregnancy (17). Moreover, we have shown that low maternal adiponectin levels in obese pregnant mice is mechanistically linked to enhanced placental function and fetal overgrowth (18), as well as the development of metabolic (20) and cardiovascular disease (19) in the offspring. Thus, understanding the molecular mechanisms governing adipose adiponectin secretion in pregnancy may help us understand the cause of impaired glucose metabolism in pregnancies among women with obesity and GDM. Furthermore, it potentially provides critical insights into the molecular mechanism linking the endocrine function of the maternal adipose tissue with placental function, fetal growth and programming of adult disease.

We explored the impact of obesity in pregnancy on adipose tissue adiponectin levels and ubiquitination, insulin signaling and ER stress using a mouse model. Specifically, in this model, obesity is induced in female mice prior to mating by feeding a diet high in saturated fat, cholesterol and simple sugars (18,31), resembling a diet common in Western societies (42). This mouse model results in fetal overgrowth associated with maternal metabolic alterations similar to those observed in obese pregnant women (14,18,31,43). Using this mouse model, we report for the first time that maternal obesity in pregnancy is associated with decreased adipose tissue adiponectin levels and increased adiponectin ubiquitination, consistent with the decreased circulating adiponectin concentrations (18,31). Moreover, our data on IRS-1 (Ser307) and Akt (Thr308) phosphorylation, together with elevated circulating insulin levels (18,31) suggest that adipose tissue in obese dams is insulin resistant. Moreover, inflammation signaling through JNK (44) and ER-stress (30) is known to phosphorylate IRS-1 at ser307. Thus, we propose that the observed adipose inflammation, and in particular JNK activation and ER stress in adipose tissue in obese mice, represents the mechanistic underpinnings of adipose tissue insulin resistance in obese pregnant mice (**Figure 5**).

To confirm the clinical relevance of our findings, we collected omental (visceral) adipose tissue at c-section from term normal BMI and obese women. We observed an increase in adiponectin ubiquitination in obese pregnant women, in agreement with our observations in our mouse model of obesity in pregnancy. However, total adipose adiponectin was not altered in obese women as compared to pregnant women with normal BMI. This suggests that the fraction of total adiponectin that is ubiquitinated is higher in adipose tissue of obese pregnant women as compared to adipose tissue of pregnant women with normal BMI. Circulating adiponectin levels were also not statistically significantly different between pregnant women with normal or obese BMI in our study, although HMW to total adiponectin ratio was trending lower in obese women (Table 1). There may be many possible explanations for this finding. First, this is a small subgroup of a larger cohort of women in which we observed the expected inverse correlation between maternal BMI and circulating adiponectin (unpublished data). Moreover, the subjects from which adipose tissue was collected for the current study were selected randomly based on BMI and not by adiponectin levels. By chance, the adiponectin levels in the obese women in this subgroup tended to be higher than in the overall cohort, and the adiponectin levels in the lean women in this subgroup tended to be lower than in the entire cohort (unpublished). Additionally, despite the high BMI, the obese pregnant women in this group may be metabolically healthy as evidenced by normal cholesterol levels (Table 1). Nevertheless, adiponectin ubiquitination was inversely correlated with circulating maternal adiponectin levels in the subset of these women with available adipose tissues. These findings suggest that although adiponectin ubiquitination plays a crucial role in maternal adiponectin secretion, additional factors may contribute to maternal circulating adiponectin levels during pregnancy.

We provide compelling evidence that insulin increases cellular adiponectin protein abundance by decreasing adiponectin ubiquitination, resulting in reduced UPS-mediated degradation using differentiated 3T3-L1 adipocytes, a widely used in vitro model of white adipocyte biology (33,45). These findings are in general agreement with literature reports showing that insulin stimulates the secretion of adiponectin in 3T3-L1 adipocytes and in non-pregnant animals and humans (21–23). Post-translational regulation of adiponectin protein abundance and secretion is established (46,47), whereas the role of the UPS in regulating adiponectin secretion is only beginning to be explored. Wang and co-workers reported that exposure of primary mouse adipocytes to 4-hydroxynonenal, a lipid peroxidation end product inducing oxidative stress, caused a decrease in adiponectin secretion by accelerating ubiquitin-proteasome degradation of the adipokine (28). Moreover, inhibition of MEK/ERK1/2 signaling accelerated UPS-mediated adiponectin degradation in cultured mouse adipocytes (27).

Obesity is associated with inflammation and ER stress in adipose tissue (29), which inhibit insulin signaling (30). We found that inflammation and ER stress inhibits the effects of insulin on adiponectin abundance, ubiquitination and degradation in differentiated 3T3-L1 adipocytes. These results suggest a role for UPS in mediating the decline in adiponectin production associated with adipose tissue inflammation and ER stress. The findings that insulin signaling inhibits adiponectin degradation support insulin’s role in regulating proteostasis (48), in part through the inhibition of the UPS (49). For example, activation of the UPS caused by insulin resistance is linked to protein degradation and skeletal muscle wasting (50). Therefore, impaired adiponectin secretion due to insulin resistance may be one of several phenotypes associated with the regulation of the UPS by insulin signaling. While it is clear that adiponectin sensitizes insulin’s actions, the role of insulin in regulating adiponectin secretion is only now being recognized. Our findings suggest that regulation of adiponectin secretion and insulin signaling are interrelated and may explain why individuals with insulin resistance exhibit low adiponectin levels.

In conclusion, we show that adiponectin protein is regulated by the UPS and that insulin inhibits the UPS leading to elevated adiponectin abundance whereas inflammation and ER stress, associated with maternal obesity accelerate adiponectin ubiquitination and degradation. This information will help us better understand the mechanisms controlling maternal insulin resistance and fetal growth in normal and complicated pregnancies and may provide a foundation for the development of strategies aimed at improving adiponectin production in pregnant women with obesity or gestational diabetes.

## Acknowledgments

The imaging experiments were performed in the Advanced Light Microscopy Core part of NeuroTechnology Center at University of Colorado Anschutz Medical Campus supported in part by Rocky Mountain Neurological Disorders Core Grant Number P30 NS048154 and by Diabetes Research Center Grant Number P30 DK116073.

## Author Contributions

I. Aye, T. Jansson, and T. Powell designed research; I. Aye, F. Rosario, O. Kristansen and A. Kramer analyzed data; I. Aye, F. Rosario, O. Kristiansen A. Kramer, and T. Michelsen performed research. I. Aye, F. Rosario, T. Michelsen, T. Jansson, and T. Powell wrote the paper. T Jansson is the guarantor of this work and, as such, had full access to all the data in the study and takes responsibility for the integrity of the data and the accuracy of the data analysis.

### Funding

The present work was supported by NIH (HD065007).

